# Reactive Bergmann glia play a central role in Spinocerebellar ataxia inflammation via the JNK pathway

**DOI:** 10.1101/2022.06.29.498121

**Authors:** Chandrakanth Reddy Edamakanti, Vishwa Mohan, Puneet Opal

## Abstract

The spinocerebellar ataxias (SCAs) are devastating neurological diseases characterized by progressive cerebellar incoordination. While neurons bear the brunt of the pathology, a growing body of evidence suggests that glial cells are also affected. It has, however, been difficult to understand the role of glia, given the diversity of subtypes, each with their individual contributions to neuronal health. Using human SCA autopsy samples we have discovered that Bergmann glia—the radial glia of the cerebellum, which form intimate functional connections with cerebellar Purkinje neurons—display inflammatory JNK-dependent c-Jun phosphorylation. This phosphorylation defines a signaling pathway not observed in other activated glial populations, providing an opportunity to specifically isolate the role of Bergmann glia in SCA inflammation. Turning to an SCA1 mouse model as a paradigmatic SCA, we demonstrate that inhibiting the JNK pathway reduces Bergmann glia inflammation accompanied by improvements in the SCA1 phenotype both behaviorally and pathologically. These findings demonstrate the causal role for Bergmann glia inflammation in SCA1 and point to a novel therapeutic strategy that could span several ataxic syndromes where Bergmann glia inflammation is a major feature.

**Significance Statement:** We have identified a Bergmann-glia specific signaling pathway that contributes to cerebellar degeneration in the spinocerebellar ataxias. This pathway is defined by activation of JNK that phosphorylates the transcription factor c-Jun leading to the release of IL-1β and potentially other cytokines from Bergmann glia. Inhibiting c-Jun phosphorylation with pharmacological JNK inhibition could serve as therapeutic approach to treating cerebellar degeneration.

## Introduction

The spinocerebellar ataxias (SCAs) are a group of autosomal dominant disorders characterized by adult onset cerebellar and brainstem degeneration. They are progressive and untreatable, and patients eventually die from respiratory complications such as aspiration and pneumonia (1-3). The more prevalent SCAs are caused by CAG trinucleotide genomic expansions. These mutations occur in the coding region of the relevant gene and therefore result in an expanded polyglutamine tract in the encoded protein. The polyglutamine ataxias include SCAs 1, 2, 3, 6, 7, and 17 and a related ataxic syndrome, dentatorubral-pallidoluysian atrophy (3, 4). Together, they account for approximately 80% of the genetically elucidated SCAs ^(5)^. The rest are caused by either microsatellite repeats or conventional mutations such as deletions or point mutations that alter the coding region of the affected genes (6).

It is not entirely clear why the SCAs display regional vulnerability as reflected in the early involvement of the cerebellum that continues to degenerate as the disease progresses. The mutant proteins themselves are ubiquitously expressed and at largely similar levels in different cell type and tissues. To understand selective vulnerability, most of the focus has been on understanding cell autonomous changes in neurons, most notably Purkinje neurons that show dystrophic changes. But other neuronal populations, particularly those of the cerebellum and brainstem, also contribute to the syndrome. These include the inferior olivary neurons of the medulla, which provide glutamatergic excitatory inputs to Purkinje cells (PCs) by their dendritic connections, termed climbing fibers, and molecular layer interneurons in the cerebellar cortex, which provide GABAergic inhibitory inputs to further sculpt PC output (7, 8). Other brainstem and cranial nerve nuclei including the vagus and hypoglossal are also involved to variable extents in the different SCAs. These studies have been performed in human autopsy samples and in genetically engineered mouse models of the disease defining the neuronal circuitry underlying the SCA syndromes (9, 10).

In addition to the neurons, however, there is also a growing awareness that non-neuronal cells such as endothelial cells and glia contribute to pathology (11-15). Pathology in these non-neuronal cells could compromise neuronal function by interfering with their normal supportive role or triggering neuroinflammation. There is considerable empiric evidence for the latter from imaging modalities such as magnetic resonance spectroscopy, and pathological analysis of patients at autopsy (16-18). The role of glia is particularly intriguing since they display alterations in gene expression which parallel those seen in neurons both based on the magnitude of alterations and longitudinal changes as the disease progresses (19, 20). Indeed, gene expression changes are seen in all the major glial populations: oligodendrocytes that ensheath neurons: astrocytes that participate in complex neuronal-glial interactions to support neurons; and microglia, the resident macrophages, which protect neurons from stress and activating inflammatory responses (21-23). Despite these findings their causal role in pathogenesis has been difficult to decipher. Key obstacles to our understanding include the complex crosstalk of signals between glia and neurons, and the remarkable diversity beyond even the basic glial classification outlined above.

The cerebellum, for instance, has at least three distinct types of astrocytes—fibrous astrocytes in the deep white matter; protoplasmic or velate astrocytes in the granular layer; and Bergmann glia (BG), regionally specialized radial astrocytes that closely align with PCs (24-26). Cerebellar microglia and oligodendrocytes also show distinct subpopulations whose diversity is still largely unexplored (27, 28). One could in fact imagine a scenario where some glia participate in a pathogenic manner while yet others provide compensatory neuroprotective signals.

Here we describe that BG are unique among the glial population of the cerebellum in that they express abundant levels of c-Jun, a prototypical member of the Jun family of nuclear factors. This transcription factor is activated by phosphorylation and heterodimerizes with other transcription factors such as Fos, ATF and CREB family members to trigger an inflammatory cascade (29). This uniqueness allowed us to intervene pharmacologically to tamp down BG-specific inflammation.

We decided to use SCA1 as a paradigm for these studies, since we and others have already established the presence of significant neurogliosis that affects BG (15, 22). SCA1 also is the most severe of the polyglutamine ataxias in terms of disease progression in humans (30, 31). Using SCA1 knock-in mice, we discovered that BG inflammation indeed plays a deleterious role in SCA, that can be thwarted by a c-Jun N-terminal kinase (JNK) inhibitory drug that prevents c-Jun activation. These results unequivocally establish the harmful role of BG inflammation in abetting cerebellar degeneration and inspire the first glia-based strategy to treat the SCAs.

## Results

### SCA patients display BG-specific phosphorylation of the transcription factor c-Jun

Bacterial lipopolysaccharide (LPS), a potent inducer of inflammation, is a major component of the outer membrane of Gram-negative bacteria. It stimulates Toll-like receptors that in turn signal through the MAP kinase receptor family of serine/threonine kinases to result in the phosphorylation of transcription factors (32). This signaling module culminates in the transcriptional expression of downstream inflammatory factors (33, 34).

We used these properties of LPS to study cerebellar inflammation in an *in vitro* system. Specifically, we treated primary mixed cerebellar cultures derived from new mouse pups to LPS added to the culture medium. In the course of studying the signaling cascades downstream of LPS we observed a robust phosphorylation of the inflammatory transcription factor c-Jun (**Supplementary Figure 1A)**. This phenomenon has been previously described (35-37). The phosphorylation of c-Jun occurs on serine 63 (c-Jun-pS63), which is known to be a serine residue targeted by JNK ^(38)^. However, c-Jun phosphorylation does not occur in all cells; it is confined to a subpopulation of glial cells that we later identified as BG based on co-staining with S100, a calcium binding protein that is solely expressed by these cells in the cerebellum (39). Total c-Jun positive cells did not change significantly compared with control cultures, indicating that c-Jun phosphorylation, and not its levels, is increased with LPS activation. We confirmed BG-specific c-Jun phosphorylation upon LPS induction *in vivo* by intraperitoneal injection of mice with LPS and staining for phosphorylated c-Jun (**Supplementary Figure 1B**).

Since gliosis is a major feature of the SCAs, we next asked whether the SCAs mirror the LPS-induced induction of c-Jun phosphorylation specifically in BG. At this juncture it is important to note that the pattern of gliosis varies in the different SCAs. For instance, in virtually all the SCAs gliosis occurs in the cerebellum; SCA3 however is an exception in that inflammation is confined to the pons ^(40, 41)^. To test c-Jun activation in different SCAs, we turned to autopsy samples from patients with SCAs. We focused on SCAs 1, 2, 3 and 7. We observed robust phosphorylation of c-Jun in BG in SCAs 1, 2, and 7, but not SCA3 (**Figure 1**). Together, these results demonstrate that BG activation as defined by c-Jun phosphorylation is a broad but not universal phenomenon across the SCAs, with the intensity of c-Jun activation corresponding to those SCAs with the most visible cerebellar inflammation.

**Figure 1:**
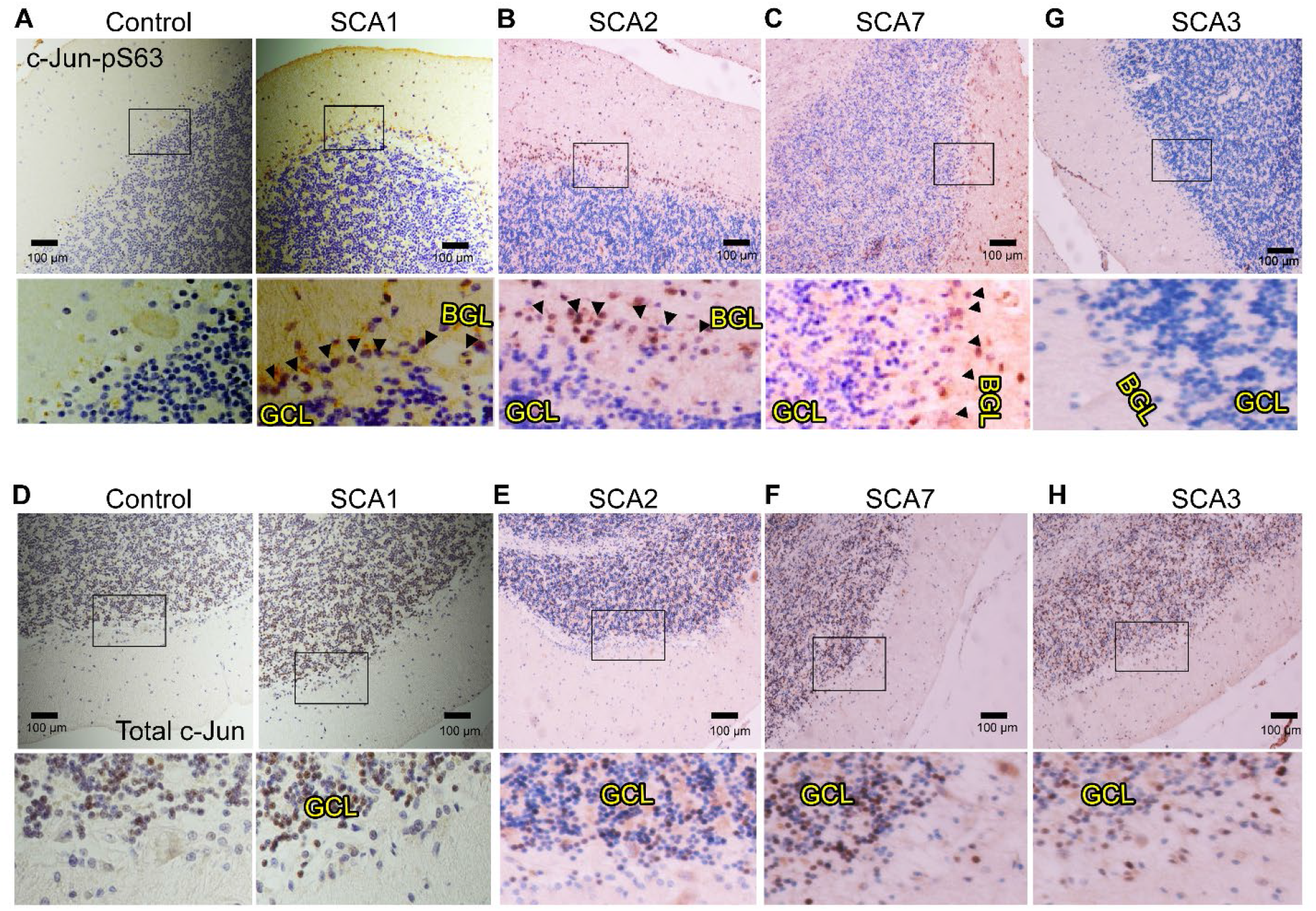
Spinocerebellar ataxia patients exhibit Bergmann glia-specific c-Jun phosphorylation. (**A-D**) HRP-DAB immunostaining of human cerebellum from patients with (**A**) SCA1 (**B**) SCA2 (**C**) SCA7 and (**G**) SCA3 using c-Jun phosphorylation (c-Jun-pS63) antibody. Black-boxed regions represent the corresponding higher-magnification images shown below each photo. (**E-H**) HRP-DAB immunostaining of human cerebellum from patients with (**D**) SCA1 (**E**) SCA2 (**F**) SCA7 and (**H**) SCA3 using total c-Jun antibody. Black-boxed regions represent the corresponding higher-magnification images shown below each photo. In all panels nuclei are counterstained with hematoxylin. Scale bar = 100μm. Representative images are shown. BGL: Bergmann Glia Layer; GCL: Granular Cell Layer. We performed staining on multiple SCA1 (n=4), SCA2 (n=3), SCA3 (n=3), and SCA7 (n=3) samples, as well as age-matched controls (n=4). Arrowheads correspond to phosphorylated c-Jun staining in BG.

### Characterizing c-Jun phosphorylation in SCA1 mice

To delve into BG inflammation in more detail, we turned to the SCA1 knock-in model. This mouse is engineered to express ATXN1 harboring 154 glutamine repeats. Even though all the functions of ATXN1 have yet to be deciphered, it appears to serve as a transcriptional regulator, whose functions and interactors are affected by the pathogenic expansion (42). This SCA1 knock-in is an extremely precise model of human disease in that mutant ATXN1 is expressed under its endogenous promoter thus mirroring the spatial and temporal pattern of ATXN1 expression. It has also been extremely well-characterized with established timelines to monitor behavioral and pathological changes spanning the mouse lifespan (43, 44). It is important to note that the polyglutamine tract in normal ATXN1 in humans is variable but does not extend beyond 40 repeats (whereas mouse ATXN1 has only 2 glutamines). The maximum glutamine repeat length described in human patients is 82 and causes a childhood onset of the disease, but this repeat must be further exaggerated to ensure a robust ataxic phenotype in the short lifespan of the mouse.

We stained for the BG-specific protein S100 along with c-Jun-pS63. The number of S100/c-Jun-pS63 double-positive cells was significantly increased in SCA1 cerebellum compared to wild-type controls (**Figure 2A-D**). Phosphorylation of c-Jun was observed as early as 16 weeks of age, a time when BG also display glial activation evidenced by GFAP staining (**Figure 2E-F**). These results extend our finding of BG-specific c-Jun activation to genetically engineered mice.

**Figure 2:**
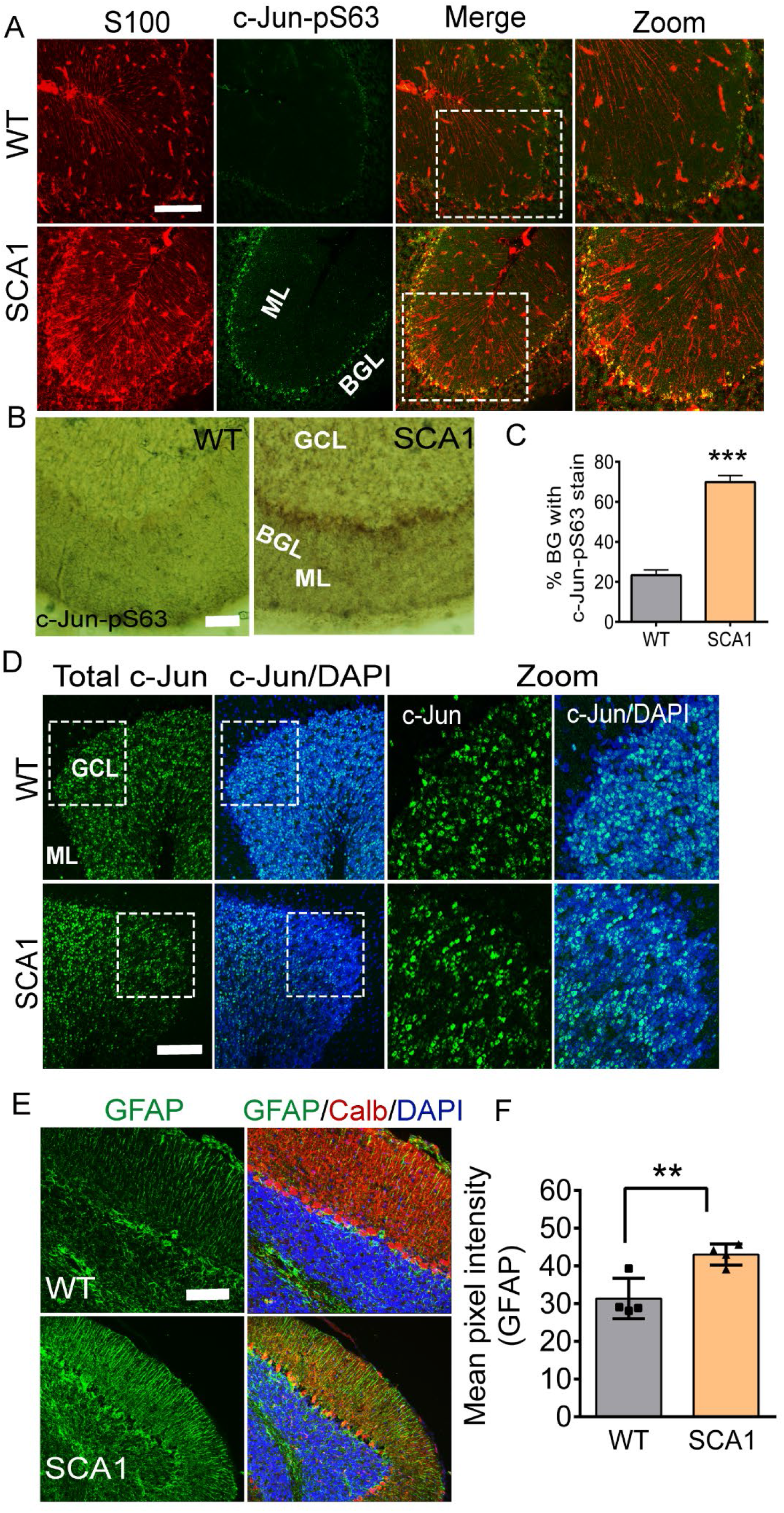
c-Jun phosphorylation in the activated Bergmann glia of SCA1 mice. (**A**) Immunostaining of 16-week-old SCA1 mouse cerebellum with Bergmann glia (BG)-specific S100 (red) along with anti-c-Jun-pS63 antibody (green). Scale bar = 100μm. White-boxed regions represent the corresponding higher-magnification images shown in the “zoom” panels. (**B**) HRP-based DAB immunostaining of SCA1 mouse cerebellum with anti-c-Jun-pS63 antibody. Scale bar = 50μm. (**C**) Quantification of percentage of BG cells positive for c-Jun-pS63 stain shown in panel A. (**D**) Immunostaining of SCA1 mouse cerebellum with total c-Jun (green). Sections are counterstained with DAPI (blue) to detect nuclei. Scale bar = 100μm. (**E**) Immunostaining of 16-week-old SCA1 mouse cerebellum with glia-specific GFAP antibody (green) along with calbindin antibody to detect Purkinje cells (red). Scale bar = 100μm. (**F**) Quantification of GFAP intensity. BGL: Bergmann Glia Layer; GCL: Granule Cell Layer; ML: Molecular Layer. Sections are stained for nuclei using DAPI (blue). n=4 mice. **P < 0.01; ***P < 0.001, 2-tailed unpaired Student’s *t* test.

We also confirmed elevated c-Jun phosphorylation specifically in BG in cerebellar dissociated cultures from SCA1 postnatal mice compared to cultures derived from wild-type mice (**Supplementary Figure 2**).

### JNK inhibitor treatment in SCA1 mice abolishes c-Jun phosphorylation and inhibits the levels of the cytokine IL-1β

Since there are well-characterized inhibitors of JNK catalytic activity, we turned to a pharmacological approach to inhibit JNK kinases. Encoded by three distinct genes, JNK kinases exist as three types: JNK 1, 2, and 3, each with further sub-types resulting from differential splicing (45). As we yet do not know which JNK subtype is responsible for c-Jun activation in BG, we used a broad JNK inhibitor, SP600125 (inhibiting JNK1 and 2 with an IC50 of 40nM, and JNK3 with an IC50 of 90nM) (46). This compound crosses the blood-brain barrier and has been used to study the role of JNK kinases in the CNS (47-49).

We treated SCA1 mice and wild-type littermates with intraperitoneal injections of SP600125 using a previously established delivery schedule (47-49). As expected, mice treated with this drug displayed a decrease in c-Jun phosphorylation in the BG layer (**Figure 3A, B and D**). This inhibition was accompanied by a reduction of the glial activation marker GFAP, demonstrating that JNK activation is required for BG activation (**Figure 3C and E**).

**Figure 3:**
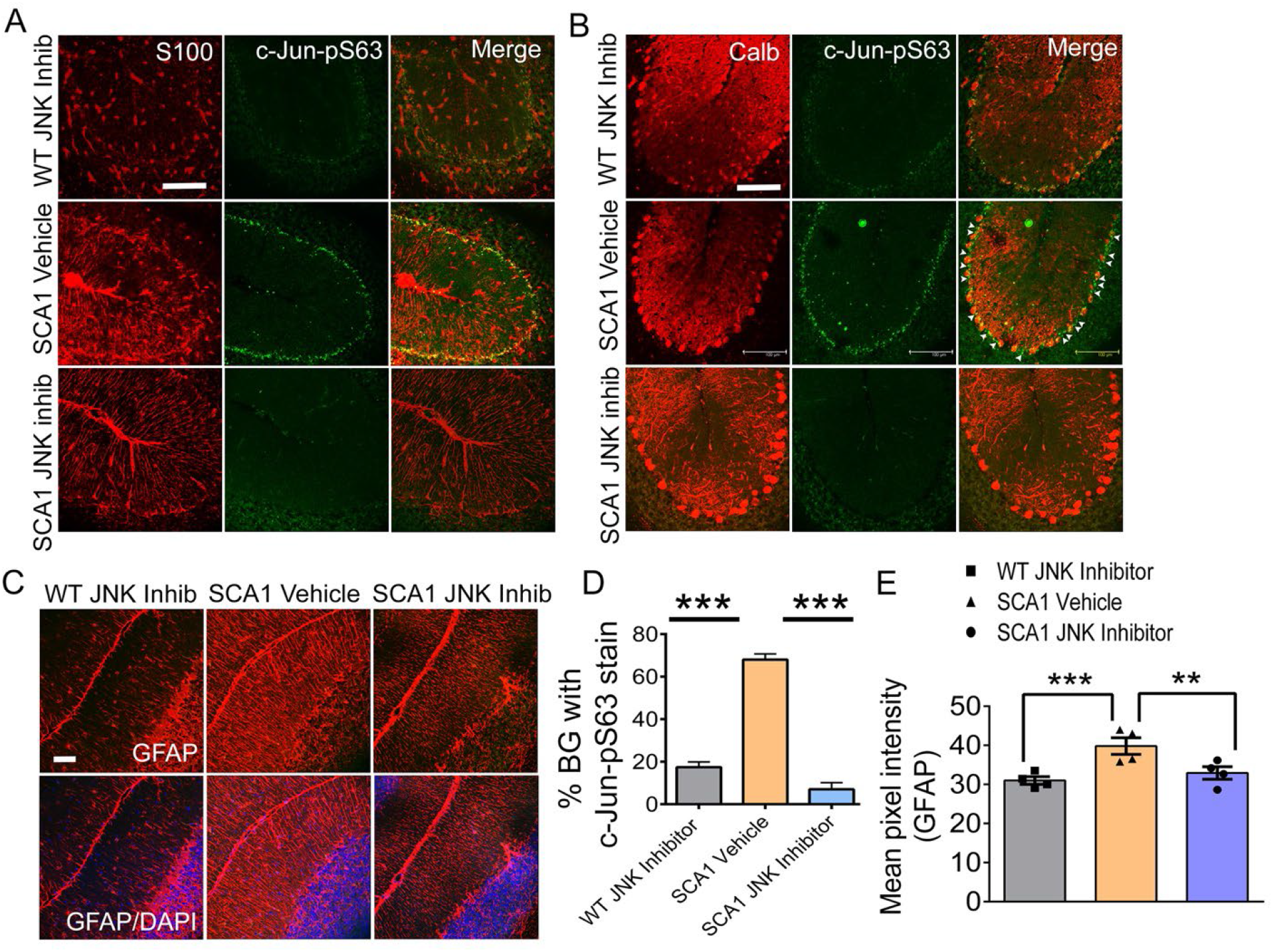
Treatment of SCA1 mice with JNK inhibitor abolishes c-Jun phosphorylation in reactive Bergmann glia. (**A**) Immunostaining of WT (JNK inhibitor) and SCA1 (Vehicle/JNK inhibitor) treated cerebellum with Bergmann glia (BG)-specific S100 (red) along with c-Jun-pS63 antibody (green). Scale bar = 100μm. (**B**) Immunostaining of WT (JNK inhibitor) and SCA1 (Vehicle/JNK inhibitor) treated cerebellum with Purkinje cell-specific calbindin (Calb) antibody along with c-Jun-pS63 antibody. C-Jun-pS63-positive BG cells (green) sit adjacent to the Purkinje cells (red). Scale bar = 100μm. (**C**) Immunostaining of WT (JNK inhibitor) and SCA1 (Vehicle/JNK inhibitor) treated cerebellum with glia-specific GFAP antibody (red). Sections were also stained for nuclei using DAPI (blue). Scale bar = 100μm. (**D**) Quantification of the percentage of BG cells positive for c-Jun-pS63, as shown in panel A. (**E**) Quantification of GFAP fluorescence intensity shown in panel C. n= 4 mice. **P < 0.01; ***P < 0.001, one-way ANOVA with Bonferroni’s multiple comparison test. Inhib =Inhibitor.

Since a major consequence of glial activation is cytokine release, we next asked whether the reduction of BG glial activation is associated with the reduction of cytokines, at least those known to play a role in neuroinflammation. We isolated RNA from experimental and control mice and performed real-time PCR (RT-PCR) to monitor the expression levels of IL-1β, IL-6, CCL2, and IL-18— four major proinflammatory molecules previously shown to be expressed in mice upon LPS treatment (50, 51). We observed a significant increase in levels of IL-1β and IL-6 in SCA1 mice compared to wild-type controls (3.5-fold and 2-fold, respectively). Of the two, the increase in IL-1β mRNA but not IL-6 was reversed by JNK inhibition (**Figure 4A**). These results point to the inflammatory factor IL-1β but not IL-6 as a BG-specific cytokine that is released upon activation.

**Figure 4:**
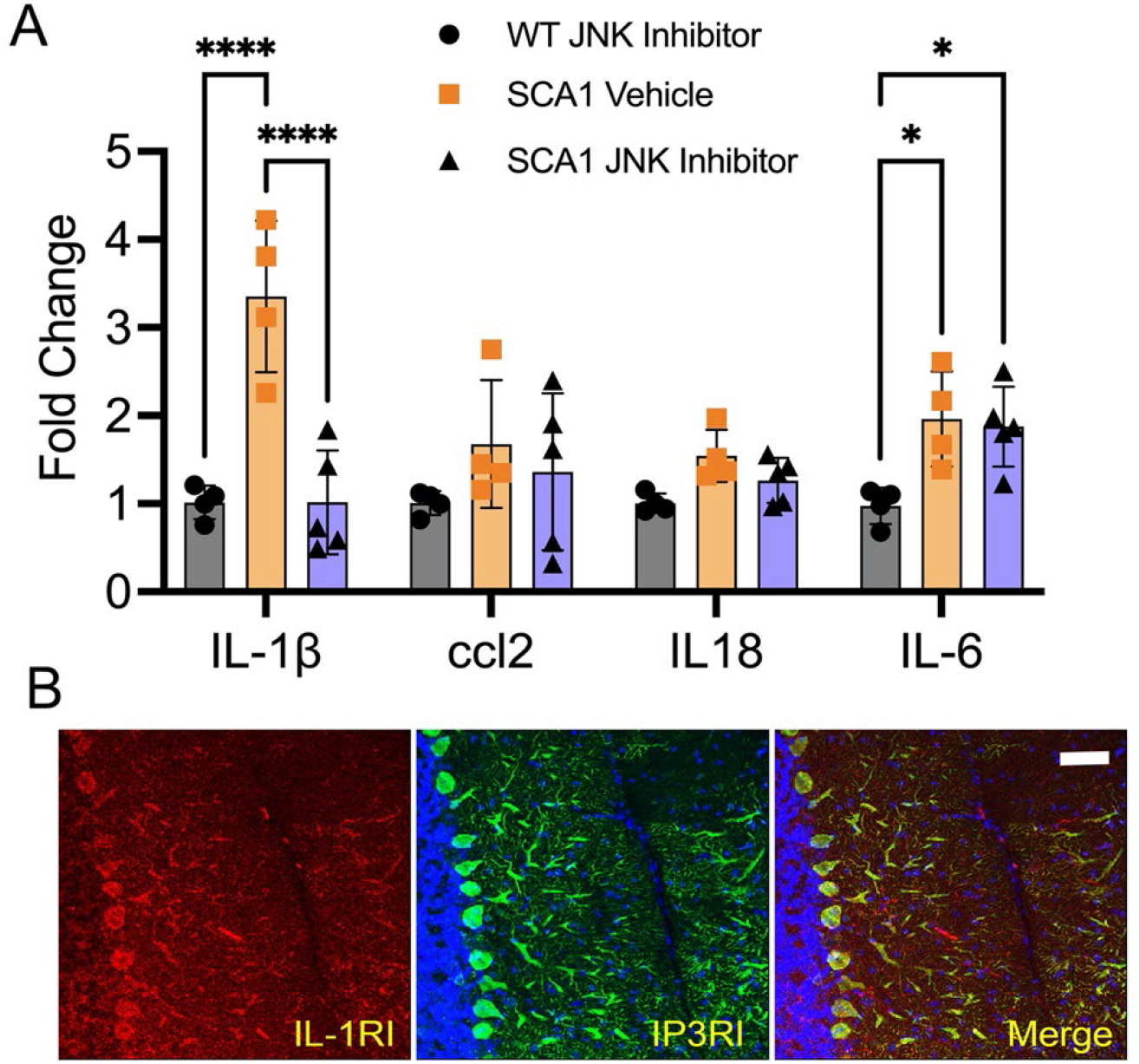
JNK/c-Jun pathway is essential for Bergmann glia-specific cytokine release in SCA1 mice. (**A**) Quantitative real-time PCR analysis of IL-1β, IL-6, CCL2, and IL-18— four major proinflammatory cytokines from SCA1/WT cerebellum either treated with vehicle or JNK inhibitor as indicated in the bar graph legend. The data were normalized to GAPDH mRNA and are represented as the fold change. (**B**) Immunostaining of WT cerebellum using IL-1 receptor (IL-1RI) antibody (red) along with Purkinje cell-specific antibody against inositol-triphosphate receptor type I (IP3RI; green). Sections were also stained for nuclei using DAPI (blue). Scale bar = 50μm. n= 3 mice. *P<0.5; ****P < 0.0001, two-way ANOVA with Tukey’s multiple comparisons test.

To identify which cells would be most affected by BG-specific release of IL-1β, we performed immunostaining of cerebellum for interleukin-1 receptor type I (IL-1RI) the predominant receptor for IL-1β ligand in the CNS. IL-1RI staining is not widespread in the cerebellum. We found the expression of this receptor in the cerebellum to be narrowly restricted to PCs (**Figure 4B**). Taken together, our results suggest a scenario wherein IL-1β (and potentially others) released by BG in SCA1 acts on PCs to cause neuronal dysfunction and death.

### JNK inhibition ameliorates the SCA1 phenotype

Neuroinflammation, including that caused by the release of cytokines such as IL-1β, can be either neuroprotective or deleterious in a broad range of pathogenic contexts including the ataxias (52, 53). To address the role of JNK activation, we studied the behavioral and pathological consequences of pharmacological inhibition of JNK in mice (**Figure 5A**). For behavioral analysis, we turned to rotarod testing, a robust measure of cerebellar motor learning, which is compromised in mouse models of the SCAs. SCA1 mice treated with JNK inhibitor showed a significant improvement in their performance. It is notable that this improvement occurred only after a month post-treatment (n=11; *P < 0.05 two months post treatment) (**Figure 5B and C**). This temporal delay is consistent with the notion that the improvement occurs because of neuroprotective effects of inhibiting inflammation. This improvement did not extend to non-cerebellar phenotypes; for instance, the weight loss in SCA1 mice– a composite sequela of neuromuscular wasting from spinal cord involvement and poor nutrition from swallowing defects– was not improved by the drug (**Figure 5D**).

**Figure 5:**
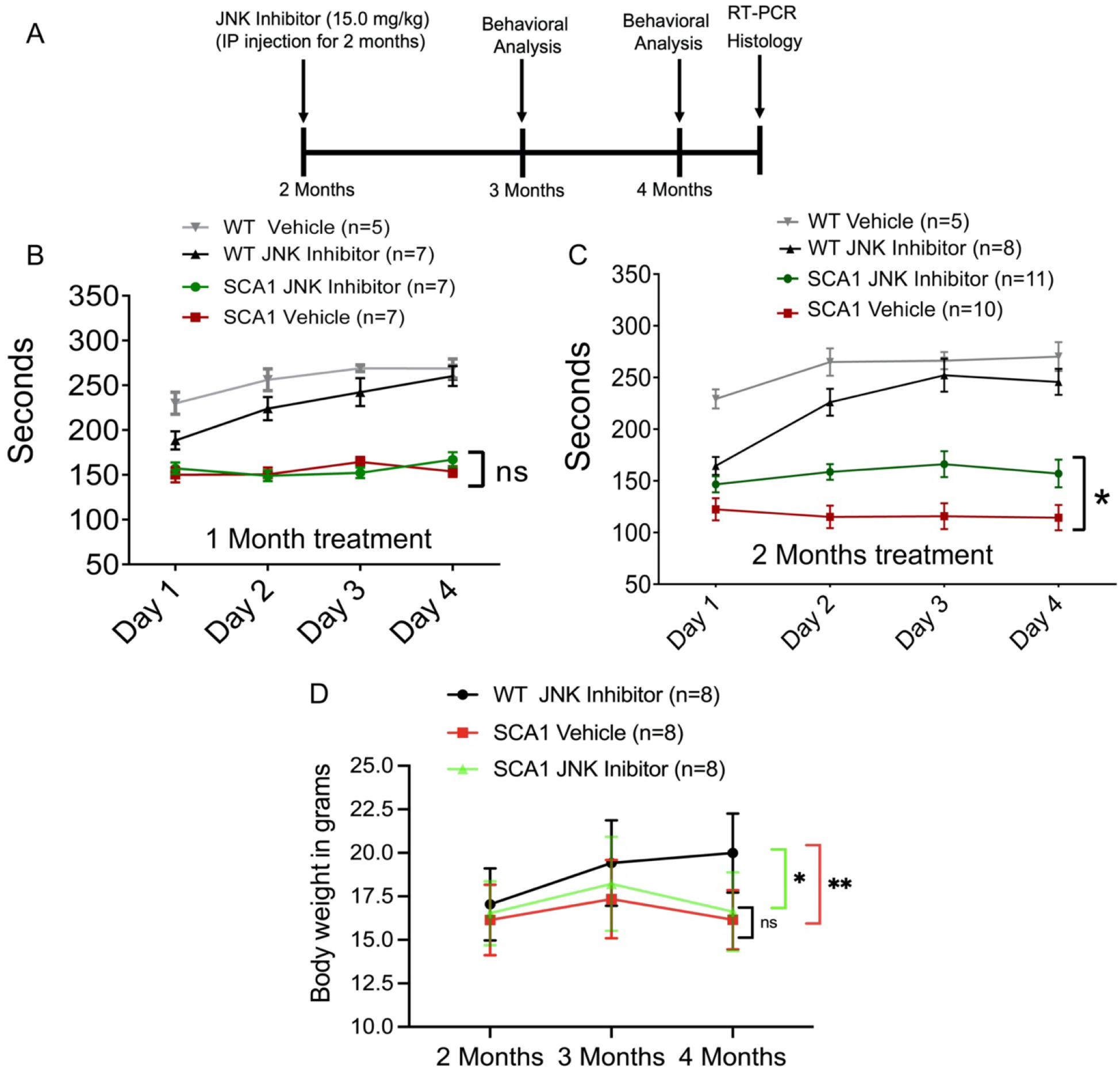
Treatment of SCA1 mice with JNK inhibitor ameliorates the motor coordination impairment. (**A**) Schematic representation of treatment and assessment course. Mice were treated with either 15 mg/kg of JNK inhibitor or vehicle (10% DMSO and 90% corn oil) intraperitoneally (IP) until 4 months of age starting from 2 months. Then mice were rested for behavioral assays followed by pathological and quantitative RT-PCR analysis. (**B and C**) Rotarod performance of mice at (**B**) three months and (**C**) four months of age. (**D**) Mouse weight before IP administration (at 2 months of age) and following administration at 3 and 4 months of age. The number of animals used is shown in the histogram legends. *P<0.05; **P < 0.01, two-way ANOVA with Bonferroni’s multiple comparisons test.

To study SCA1 cerebellar pathology, we performed experiments to address the health of PCs and their connections (54-56). Staining for calbindin, a standard marker for PCs, we observed a significant increase in the thickness of the cerebellar molecular layer in SCA1 mice treated with the JNK inhibitor compared to mice treated with vehicle control (**Figure 6A-B**).

**Figure 6:**
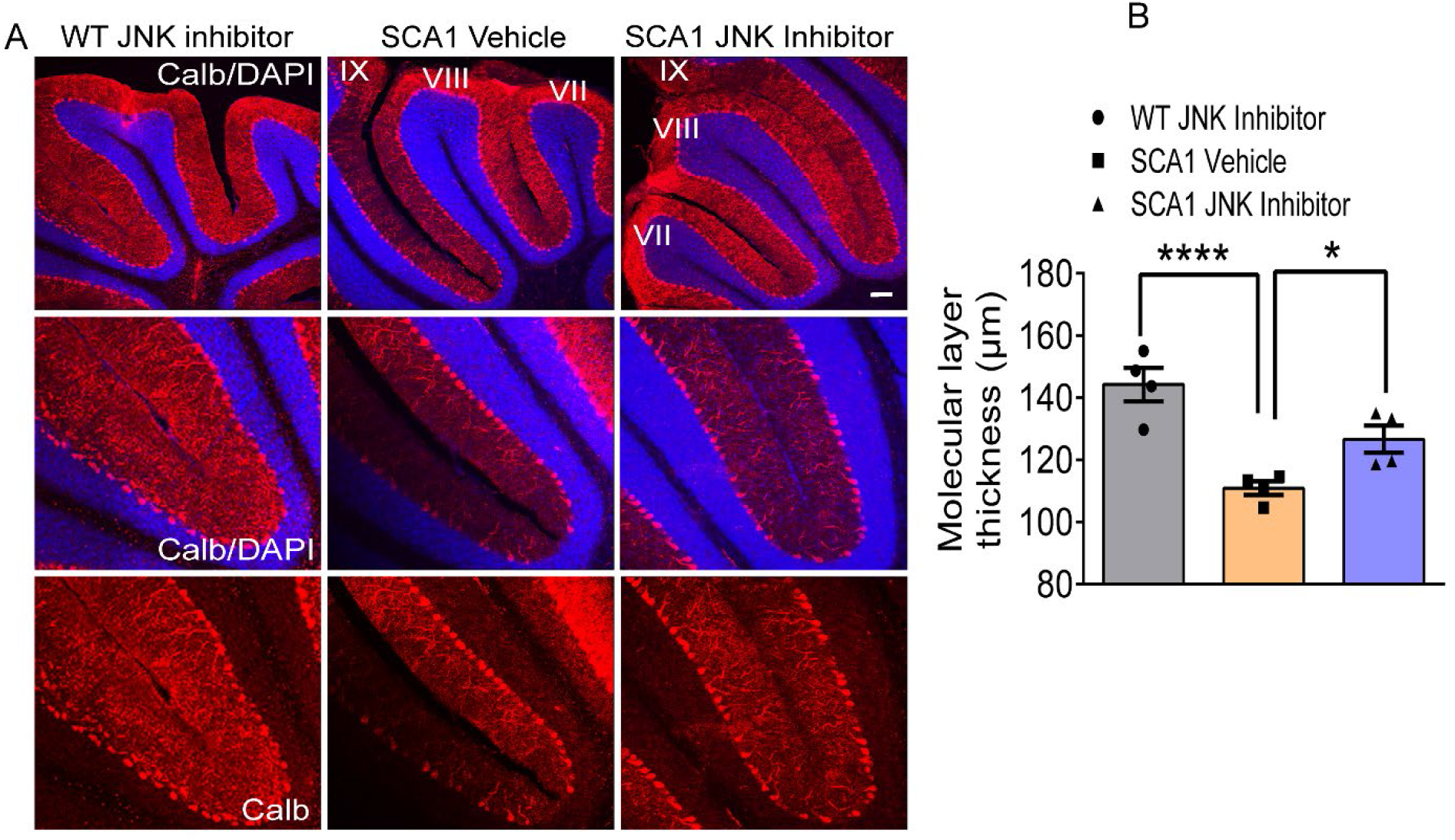
JNK inhibitor treatment improves Purkinje cell pathology in SCA1 mice. (**A**) Cerebellar slices from 4-month-old mice treated with either JNK inhibitor or vehicle were stained with calbindin antibody specific for Purkinje cells (red). Images are taken from same lobules (VII, VIII and IX) in each condition. The corresponding higher-magnification images (20X) shown below each panel. Sections were also stained for nuclei using DAPI (blue). Scale bar = 100μm. (**B**) Quantification of molecular layer thickness (Red). n= 4 mice. *P<0.05; ****P < 0.0001, one-way ANOVA with Bonferroni’s multiple comparison test.

Thus, taken together our results point to a model where BG-specific inflammation, mediated by JNK kinase, results in the release of cytokines such as IL-1β that is deleterious to PCs. This process is ameliorated by JNK kinase inhibition, which in turn improves the SCA1 phenotype (**Figure 7**).

**Figure 7:**
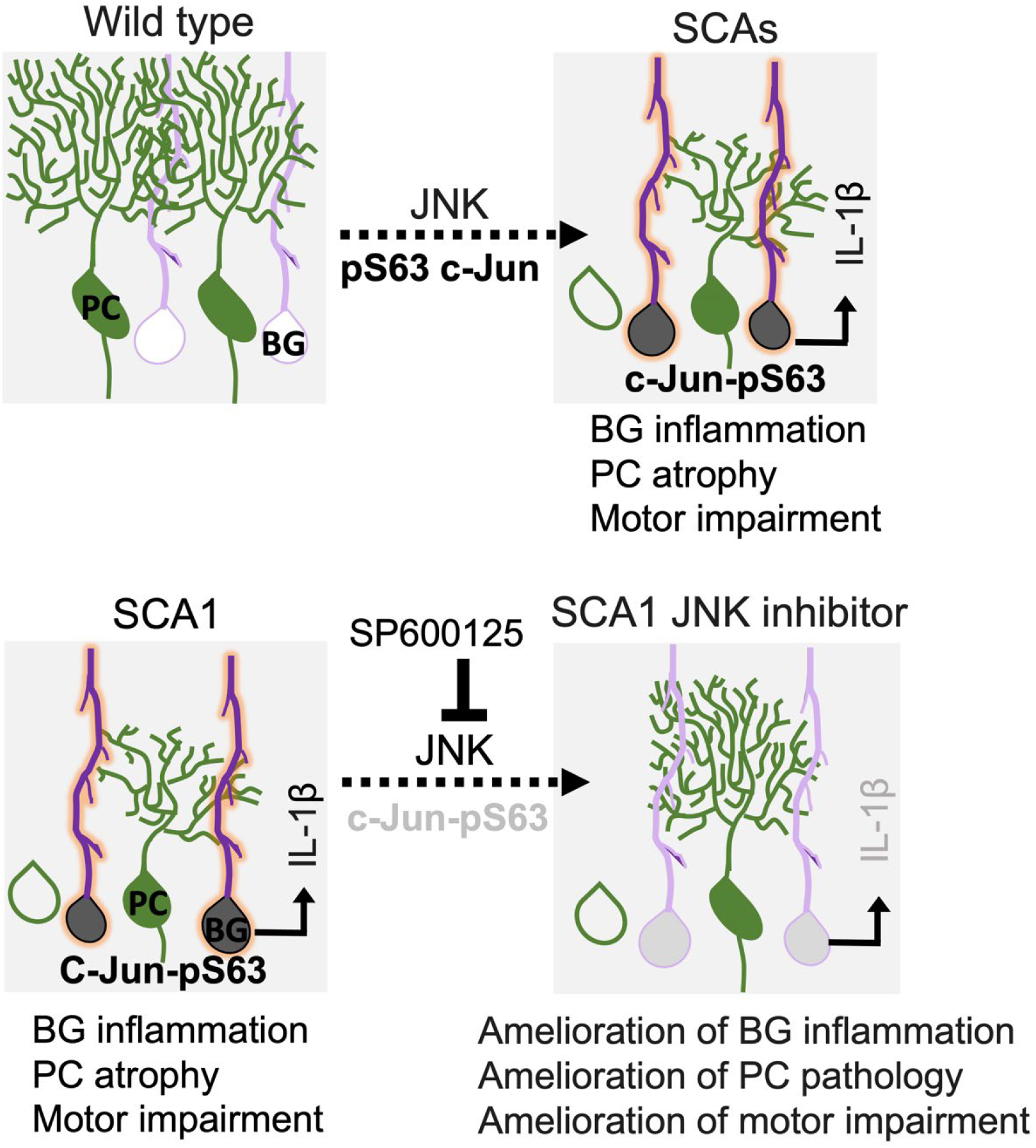
Model of targeting Bergmann glia activation to combat SCAs. **Top panel:** SCA patients and mouse cerebellums exhibit Bergmann glia (BG)-specific JNK dependent c-Jun phosphorylation (black nuclei); the time when BG are known to have a reactive state is marked by enhanced GFAP intensity (red processes). These reactive BG release the enhanced proinflammatory cytokine IL-1β into the cerebellum in a JNK-dependent manner. **Bottom panel:** Treatment of SCA1 mice with JNK inhibitor SP600125 abolishes the c-Jun phosphorylation in BG (light gray nuclei) and thereby tamps down the reactive GFAP staining and cytokine IL-1β release in the cerebellum. These changes in inflammation lead to a decrease in Purkinje cell pathology (green) and a rescue of motor deficits.

## Discussion

Neuroinflammation is a complex process reflecting an aggregate of interactions between glia, neurons, and the microvasculature. Some of these interactions are homeostatic and designed to be neuroprotective; others on the other hand are deleterious and contribute to pathogenicity (57, 58). Dissecting and identifying these complex pathways in a cell type-specific manner will be crucial to finding therapies for patients suffering from neurodegeneration.

Here we describe a signaling pathway in SCAs that is defined by BG-specific phosphorylation of c-Jun. BG are particularly well-placed to play a role in Purkinje cell dysfunction and degeneration given their location and interconnections with PCs in the cerebellar cortex; moreover they outnumber PCs 8:1 and they almost completely blanket the dendritic neuropil of PCs with their fine protrusions (24, 59, 60). The fortuitous observation of c-Jun phosphorylation in activated BG allowed us to test the relevance of BG inflammation in the context of the SCAs. Using SCA1 as a model, we discovered that BG inflammation can be tamped down by inhibiting JNK proteins that are responsible for catalyzing this signaling pathway; this treatment in turn substantially ameliorates the disease.

As we demonstrate here, one likely mechanism for downstream toxicity of BG activation is the release of cytokines, such as IL-1β, under the control of JNK signaling. Indeed, direct injection of IL-1β into the cerebellum of wild-type mice is sufficient to induce Purkinje cell pathology and cerebellar ataxia (61). But cytokine release need not be the only pathological event triggered by BG activation. BG inflammation could, for instance, also affect the normal housekeeping functions critical for maintaining neuronal health. For instance, BG are known to express the glutamate transporter EAAT2 (GLAST), which is responsible for the active reuptake of glutamate at excitatory synapses; this in turn modulates neurotransmission and prevents excitotoxicity. BG also express potassium Kir 4.1 channels, by which they regulate extracellular potassium levels in the vicinity of PCs and thus further fine-tune synaptic activity (22, 62). Perturbation of some of these normal homeostatic functions has already been hinted at in the SCAs; in conditional mouse models of SCA7, for example, BG-specific expression of mutant ATXN7 is sufficient to cause non-cell-autonomous PC degeneration by reducing the protein GLAST and causing morphological consequences of excitotoxicity (63). While similar conditional studies have yet to be performed for any of the other SCAs, SCA1 mice show a reduction in the number of BG (64), with individual glia expressing less GLAST (65). Regardless, the importance of BG to PC function has been most vividly demonstrated by optogenetic manipulation of BG, where BG inactivation leads to virtually instantaneous alterations of PC firing and subsequent cerebellar behavioral deficits (66, 67).

Our findings raise two interesting questions. First, why do BG specifically show c-Jun-dependent activation? The reason could lie in their distinct developmental origin—BG arise from a stem cell niche in the rhombic lip whereas other cerebellar astrocytes arise from stem cells derived from the cerebellar ventricular zone (24, 25). Alternatively, the role of BG in sustaining PC health may require distinct signaling pathways not shared by other glia. The second question is mechanistic, and that is: what are the proximate triggers for BG inflammation? Given that BG activation occurs even when the mutant protein is only expressed in Purkinje neurons (15). We suspect that dysfunctional PCs, perhaps in conjunction with microglia, release signals that trigger BG activation. These signals are likely to be cytokines and chemokines, since these factors have been shown to activate astrocytes in other disease contexts (68). One could thus envisage a scenario where PCs trigger BG activation, which in turn exacerbates PC dysfunction, resulting in a vicious feed-forward cycle of neurodegeneration.

An important translational contribution of this study is that it inspires a novel and eminently feasible treatment strategy for SCAs, namely, the use of JNK kinase inhibitors. While we have used a broad-specificity JNK inhibitor in these proof-of-principle studies, it would be important to determine which of the three JNK isoforms need to be targeted in order to reduce the potential side effects. It is also possible that interfering with downstream targets of c-Jun activation such as decreasing the levels or activity of IL-1β could also prove therapeutic, providing yet additional treatment avenues. We should also emphasize that while we have focused on the polyglutamine SCAs, we suspect that JNK-dependent BG activation is likely a widespread phenomenon of other cerebellar syndromes both familial and sporadic ^(69-71)^. It will be important to determine the extent to which cerebellar gliosis occurs in diverse ataxias using magnetic resonance studies or autopsy evaluations. These studies could provide the impetus for broadening this glial-based therapeutic approach.

## Materials and Methods Mouse lines

The *Sca1*^*154Q/2Q*^ line was generated by inserting a small conserved region containing 154 CAG repeats of the human sequence into the mouse *ATXN1* locus (43). Animal experiments were performed in compliance with the National Institutes of Health’s Guide for the Care and Use of Laboratory Animals and the Northwestern University Institutional Animal Care and Use Committee.

### Primary cultures of cerebellar neurons/glial cells

Neuronal/glial cerebellar cultures were derived from mice using an established protocol (38, 72). Isoflurane anesthetized mice were sacrificed by decapitation at post-natal day 4 (P4). The cerebella were dissected away from the meninges and choroid plexus. Minced cerebellar tissue was trypsinized for 15 min at 37 °C and then triturated in Hank’s Balanced Salt Solution containing 10 U/mL DNAse I (Roche Diagnostics). The cells were centrifuged at 2,000 rpm for 7 min and resuspended in Neurobasal media (Sigma) containing 4 mM glutamine, 10% FBS, 100 U/mL Penicillin/streptomycin, and 25 mM KCl (Sigma-Aldrich). After counting, 7.5 × 10^5^ cells were plated on precoated poly-D-lysine glass coverslips in 24-well plates. Cultures were maintained at 37 °C, 5% CO2, and media were changed every 2 days. On day 6 in culture, the cells were treated with LPS (Sigma #L2630) at 100 ng/mL concentration for 3 h. They were then fixed in 4% paraformaldehyde for immunohistochemical staining.

*In vivo* LPS treatment in wild-type mice was performed as previously described (38). Briefly, LPS in PBS was administered intraperitoneally at a dose of 750μg/kg for seven consecutive days. The control mice received the vehicle PBS alone. After seven days of injections, mice were sacrificed for the immunohistochemical analysis.

### Human brain immunohistochemistry

We obtained SCA autopsy samples: four SCA1, three SCA2, three SCA3, three SCA7, and four age-matched controls (from Arnulf Koeppen and Laura Ranum, with approval from their respective institutional review boards at the Veterans Affairs Medical Center, Albany, New York, and the University of Florida). Post-mortem cerebellar tissue from SCA patients was mounted in paraffin blocks, and 5μm-thick slices were cut from each paraffin block, processed for HRP-DAB staining, and counterstained with hematoxylin. Antigen retrieval and antibody staining was optimized at the Northwestern University Pathology Core.

### Experimental injections with pharmacological agents

The JNK inhibitor SP600125 (#HY-12041, MedChemExpress) was dissolved in 10% DMSO and 90% corn oil. It was injected intraperitoneally on an alternate day schedule at a dose of 15mg/kg starting when mice were two months of age and continuing for two months. Control mice were treated with vehicle alone. The mice were then evaluated behaviorally and pathologically in a blinded fashion. Since SCA mice do not display sex-based differences in their cerebellar phenotype, the read-outs from males and females were pooled before statistical analysis.

### Rotarod assays

Rotarod testing was performed by placing mice on a motorized rotating rod that accelerates linearly from 4 to 40 rotations per minute over a maximum duration of 5 minutes (Ugo Basile, Comerio, Italy) (14). The time it takes for a mouse to fall off was recorded. If mice passively clung to the rod for two consecutive rotations, that was also counted as a fall. Mice were subjected to four trials per day for four consecutive days. To ensure enough recovery time between trials, animals were given 10-15 min rest between the end of a trial and the subsequent trial.

### Pathological assays/immunohistochemistry

Mice were sacrificed by deep anesthesia (isoflurane) and transcardiac perfusion (first with PBS and then with 4% Paraformaldehyde in PBS). The brains were dissected from the cranium and post-fixed with 4% paraformaldehyde in PBS in an overnight incubation at 4 °C. They were subsequently equilibrated in a 10-30% sucrose gradient and embedded in optimal cutting temperature medium. The cerebella were sliced into 30 μm-thick sections with a cryostat (Micron M505, Thermo Fisher Scientific) or Vibratome (Leica VT1000 S).

Immunohistochemistry was then performed either by immunofluorescence or horseradish peroxidase (HRP)-based 3,3′-diaminobenzidine (DAB) detection.

For immunofluorescence, the sections were permeabilized and blocked with 10% normal goat serum and 0.25% Triton X-100 in 1x Tris-buffered saline for 1 h, after which the sections were incubated with primary antibodies (diluted in 1% BSA) overnight at 4 °C. The following day, the sections were washed three times with PBS, then incubated with fluorescently tagged secondary antibodies for 2 h at room temperature in the dark. Finally, the sections were washed three times with TBS (adding DAPI into the last wash) and mounted onto glass slides using Mowiol 4-88 (Sigma-Aldrich). The sections were imaged using a CTR6500 confocal microscope equipped with Leica LAS AF software (Leica, Buffalo Grove, IL).

For HRP-based DAB staining, the sections were processed for antigen retrieval using citrate-based buffer (pH 6.0) (Abcam #ab93678) and quenched for endogenous peroxidase activity by treating with 3% H_2_O_2_. The sections were then blocked in 10% normal blocking serum for 20 min, washed in PBS, and then incubated with primary antibody (diluted in 1% BSA) for 1 h. Sections were washed with PBS and incubated with biotinylated secondary antibody (rabbit IgG VECTASTAIN #PK-6101 or mouse IgG VECTASTAIN #PK-4002) for 30 min. After a wash with PBS, the sections were incubated with VECTASTAIN elite ABC reagent for 30 min followed by incubation with peroxidase substrate solution (VECTOR #SK-4100) for 2-10 min at room temperature until the desired brown color developed. Immediately, the slides were rinsed under tap water for 5 min. Slides were mounted using aqueous mounting medium (VectaMount AQ #H-5501).

### Quantitative real-time PCR (RT-PCR)

Mice were sacrificed by deep anesthesia (isoflurane) followed by decapitation. The cerebellar tissue was dissected from the cranium. Cerebellar RNA was extracted using an RNeasy Plus Universal mini kit (Qiagen #73404) that was then used to generate cDNA using a reverse-transcription kit (Biorad #1708840). Quantitative PCR was subsequently performed using TaqMan probes with iTaq Universal Probe Supermix on a CFX96 Real-Time thermocyler (Biorad C1000 Touch). For each sample, relative levels of target gene transcript were calculated as the ratio of Ct value of target gene (experimental to control sample) normalized to similarly derived GAPDH ratios.

The probes used were as follows: *IL-1b:* Catalog #4331182, ID: Mm00434228_m1. *CCL2:* Catalog #4331182, ID: Mm00441242_m1. *IL-18:* Catalog #4331182, ID: Mm00434225_m1. *IL-6:* Catalog #331182, ID: Mm00446190_m1. *GAPDH:* Catalog *#*4352661, Mm99999915_g1. All probes were fluorescein amidite-labeled.

## Antibodies

The following primary antibodies were used: rabbit anti-c-Jun mAb (# 9165 Clone 60A8, Cell Signaling), rabbit anti-phospho-c-Jun (Ser63) II (#9261, Cell Signaling), mouse anti-GFAP mAb (#MCA-5C10, EnCor Biotechnology Inc), rabbit anti-GFAP (# Z0334, Dako), mouse anti-IL-1RI (#AF771, R&D Systems), rabbit anti-IP3R-I (#PA1-901, Thermo Fisher), mouse anti-S100B mAb (#S2532, Sigma-Aldrich).

## Microscopy and image analyses

Nikon Eclipse TE2000-E fluorescence microscopes equipped with Intensilight C-HGFI (Nikon Inc., Melville, NY, USA) were used. Epifluorescence images were acquired using a Digital Sight DS-Qi1MC CCD camera (Nikon Inc., Melville, NY, USA), and light images were acquired using a Ds-Fi1 camera (Nikon Inc., Melville, NY, USA). Confocal images were collected using Lieca TCS SP5 confocal microscopes (Leica Inc., Bensheim, Germany) and used to acquire low-and high-magnification images of fluorescent samples. To allow for the comparative quantification of fluorescence intensity between samples, all images were acquired with standardized settings and laser power. Maximum intensity projections for individual channels were generated and the fluorescence intensity of these projections were then measured using ImageJ (NIH, Bethesda, MD, USA). Mean intensity results of 3-5 images were plotted as histograms.

## Statistical analysis

We performed all statistical tests using GraphPad Prism 4.0 (GraphPad Software). Data is presented as mean + SEM The level of significance was set at P values less than 0.05. Two tailed t-tests were used for comparison of the two data sets while two-way ANOVA and one-way ANOVA followed by Bonferroni correction were used for experiments with three or more data sets. Molecular and biochemical analyses were performed using a minimum of three biological replicates per condition.

## Study approval

All animal experiments were performed in compliance with the NIH’s Guide for the Care and Use of Laboratory Animals (National Academies Press, 2011) and were reviewed and approved by the Northwestern University IACUC.

## Author Contributions

Design and conceptualization of study: CRE and PO; Methodology and investigation: CRE, VM. Data curation: CRE, VM and PO. Writing, reviewing and editing the draft: CRE and PO.

## Acknowledgments

P.O. received support from the NIH (1R01NS082351). He has received funding from the following sources for clinical trials: Biohaven Pharmaceuticals, NIH U01NS104326 (site PI), and the National Ataxia Foundation (CRC-SCA natural history study). C.R.E received support from a young investigator award from the National Ataxia Foundation, and from a Dixon Translational Research Grant award from Northwestern University.

We thank members of the Opal lab for their suggestions throughout the course of the project, Melissa Stauffer for thoughtful comments on the manuscript, Abigail Brown for help with mouse husbandry and genotyping, and Arnulf H. Koeppen and Laura Ranum for providing SCA brain material. Autopsies were performed through a national tissue donation program supported by the National Ataxia Foundation. We thank Bella Shmaltsuyeva and the Pathology Core Facility of the Robert H. Lurie Comprehensive Cancer Center, Northwestern University, for helping with human brain immunohistochemistry.

