## Supplemental figures 1-2 for "Reactive Bergmann glia play a central role in Spinocerebellar ataxia inflammation via the JNK pathway"

### Supplementary Figures

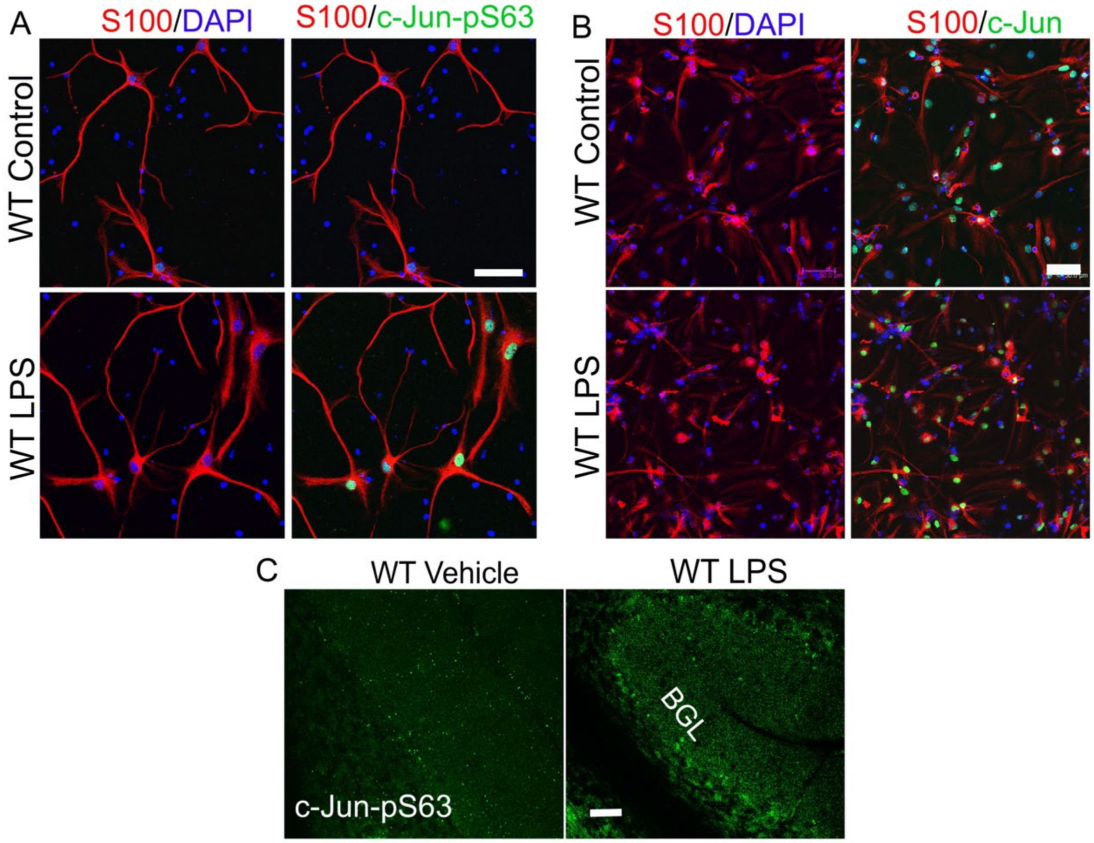


**Figure S1: Lipopolysaccharide (LPS) induces c-Jun phosphorylation in Bergmann glia *in vitro* and *in vivo.***

(**A**) Immunostaining of S100 (red) with c-Jun-pS63 (green) in DIV6 (days *in vitro* 6) neuronal/glial cerebellar cultures generated from P4 mice and treated with PBS (control) or LPS (100 ng/mL). (**B**) Immunostaining of S100 (red) with total c-Jun (green). Slides were stained for nuclei using DAPI that labeled all the cells including glia in this mixed population. Scale bars = 50µm. (**C**) Immunostaining of the cerebellum for c-Jun-pS63 antibody (green) in wild-type mice treated with LPS (750µg/kg) or vehicle (PBS) by intraperitoneal injection daily for 5 days. Scale bar = 100µm

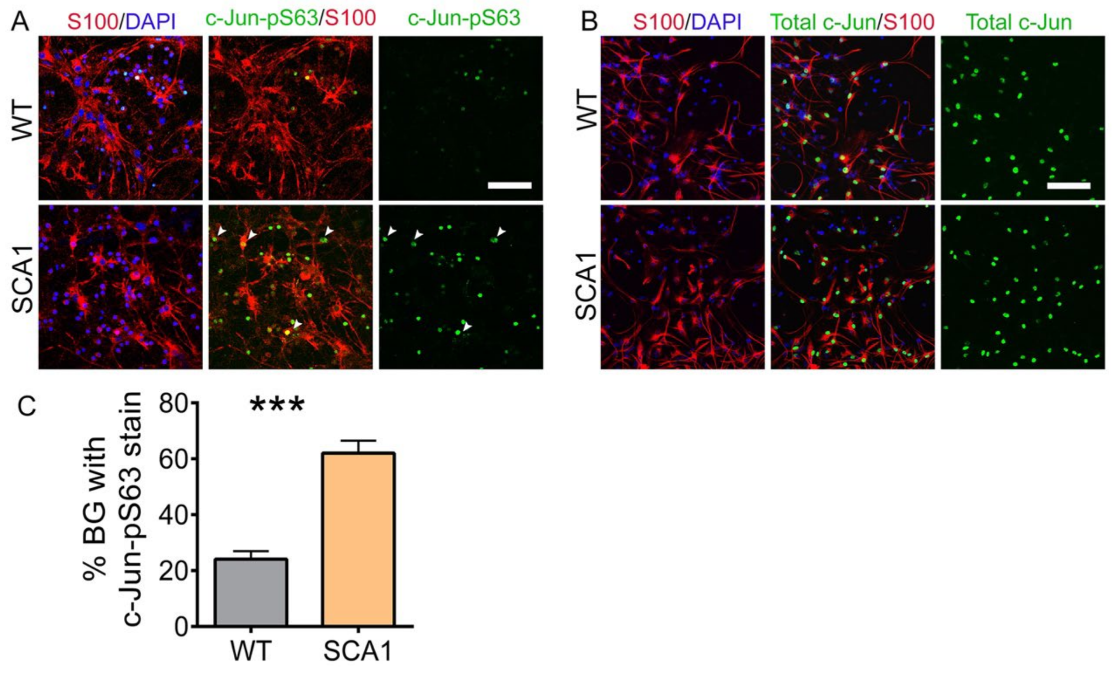


**Figure S2: *In vitro* isolated Bergmann glial cultures exhibit enhanced c-Jun phosphorylation.**

(**A-B**) DIV6 neuronal/glial cerebellar cultures generated from P4 SCA1 or wild-type mice and immunostained with S100 (red) either with (**A**) c-Jun-pS63 (left panels, green) or (**B**) total c-Jun (right panels, green). White arrowheads indicate examples of S100/c-Jun-pS63 double positive cells. Scale bar = 100µm. (**C**) Quantification of S100/c-Jun-pS63 double positives shown in panels A. n=3 individual cultures. ***P < 0.001.
